# Ultrasonic Neuromodulation Causes Widespread Cortical Activation via an Indirect Auditory Mechanism

**DOI:** 10.1101/234211

**Authors:** Tomokazu Sato, Mikhail G. Shapiro, Doris Y. Tsao

**Affiliations:** Division of Biology and Biological Engineering; Division of Chemistry and Chemical Engineering; Howard Hughes Medical Institute California Institute of Technology 1200 E. California Blvd Pasadena, CA, USA 91125

## Abstract

Ultrasound has received widespread attention as an emerging technology for targeted, non-invasive neuromodulation based on its ability to evoke electrophysiological and motor responses in animals. However, little is known about the spatiotemporal pattern of ultrasound-induced brain activity that could drive these responses. Here, we address this question by combining focused ultrasound with wide-field optical imaging of calcium signals in transgenic mice. Surprisingly, we find cortical activity patterns consistent with indirect activation of auditory pathways rather than direct neuromodulation at the ultrasound focus. Ultrasound-induced activity is similar to that evoked by audible sound. Furthermore, both ultrasound and audible sound elicit motor responses consistent with a startle reflex, with both responses reduced by chemical deafening. These findings reveal an indirect auditory mechanism for ultrasound-induced cortical activity and movement requiring careful consideration in future development of ultrasonic neuromodulation as a tool in neuroscience research.

## Introduction

The use of ultrasound to elicit targeted changes in neural activity has been the focus of intense interest in the neuroscience community due to its potential as a noninvasive technique with the ability to target deep-brain regions with millimeter precision (Landhuis, 2017). Multiple studies in rodents and other organisms have documented the ability of ultrasonic neuromodulation (UNM), applied transcranially in wild-type animals, to elicit motor responses (Tufail et al., 2010, Yoo et al., 2011, Kim et al., 2014, King et al., 2013, King et al., 2014, Ye et al., 2016, Younan et al., 2013, Mehic et al., 2014, Kamimura et al., 2016) or sensory effects (Legon et al., 2014, Lee et al., 2015, Lee et al., 2016a, Lee et al., 2016b). However, the neural signaling pathways and mechanisms underlying these responses are currently unknown, confounding the interpretation of UNM experiments in basic neuroscience research and the translation of this technique towards clinical applications. In particular, no study has analyzed cortex-wide responses to UNM to examine the neural circuits activated or inhibited by this technique, and how these circuits connect to motor behavior.

Here we address this question by imaging UNM-evoked cortical responses in mice using wide-field fluorescence microscopy. Although it is limited to monitoring the cortex, fluorescent imaging has several advantages as a readout for UNM effects compared to electrical, hemodynamic or metabolic methods. Intracranial electrical recordings are limited in the number of regions that can be sampled at the same time and the potential for artifacts due to the mechanical mismatch between electrodes and tissue, while most extracranial EEG methods have limited ability to spatially localize the sources of recorded events. At the same time, hemodynamic and metabolic techniques, such as fMRI and PET, may be confounded by ultrasonic effects on the vasculature or metabolism in addition to neural activity (Nonogaki et al., 2013, Morishita et al., 2014, Nonogaki et al., 2016, Bonow et al., 2016). By contrast, wide-field calcium imaging provides a direct readout of neuronal activation across multiple regions of the brain with relatively high spatiotemporal resolution, facilitating quantitative assessment of neuromodulation-evoked activity patterns.

Using this technique we find, surprisingly, that applying ultrasound to the visual cortex elicits cortical responses with spatial and temporal dynamics very similar to external audible sound. Moreover, both UNM and audible sound elicit motor responses consistent with a startle reflex, which are reduced with chemical deafening. These results suggest that, in addition to potentially direct neuromodulation, focused ultrasound produces secondary mechanical effects that activate auditory pathways, leading to motor responses. Together with the companion study by Guo et al. (https://doi.org/10.1101/233189) demonstrating auditory effects of UNM in guinea pigs using electrical recordings and surgical deafening, this work suggests that previous UNM studies may require re-interpretation, and that further technical developments are needed to advance UNM as a spatially precise modality for noninvasive modulation of neural circuits.

## RESULTS

### Focused ultrasound produces broad cortical activation, starting with auditory cortex

To visualize cortical responses to ultrasound, we performed simultaneous UNM and wide-field cortical imaging in transgenic Thy1-GCaMP6s mice (Dana et al., 2014) expressing the fluorescent calcium indicator GCaMP6s (Chen et al., 2013b). The mice were prepared with thinned skulls for optical access, and positioned using a surgically-implanted head-restraint bar so as to enable the imaging of the dorsal cortex while applying ultrasound (Fig. 1A). Anatomical landmarks such as Bregma and Lambda sutures, as well as large blood vessels, can be seen in the raw fluorescence image (Fig. 1B). Our ultrasound parameters were similar to those used in previous UNM studies in mice (Tufail et al., 2010, King et al., 2013, Mehic et al., 2014), with an ultrasound frequency of 500 kHz, a pulse repetition frequency of 1,500 Hz, pulse duration of 200 μs, and a total of 120 pulses per stimulation (lasting 80 ms) at intensities ranging from 0.034 W/cm^2^ to 4.2 W/cm^2^ I_SPTA_. The ultrasound was focused on the posterior portion of the visual cortex, a focus identified in multiple previous UNM studies as resulting in robust movement effects (Younan et al., 2013, Mehic et al., 2014, Ye et al., 2016, Kamimura et al., 2016). In addition, the visual cortex provides a well-known anatomical location, simple verification using light flash stimuli, and distinction from motor and auditory cortical regions. At this location, the ultrasound focus, with a half-maximal intensity diameter of 4.4 mm, lies within a single hemisphere, removed from the lateral edges of the skull, and has little overlap with other sensory cortical areas (Fig. 1C).

**Figure 1.**
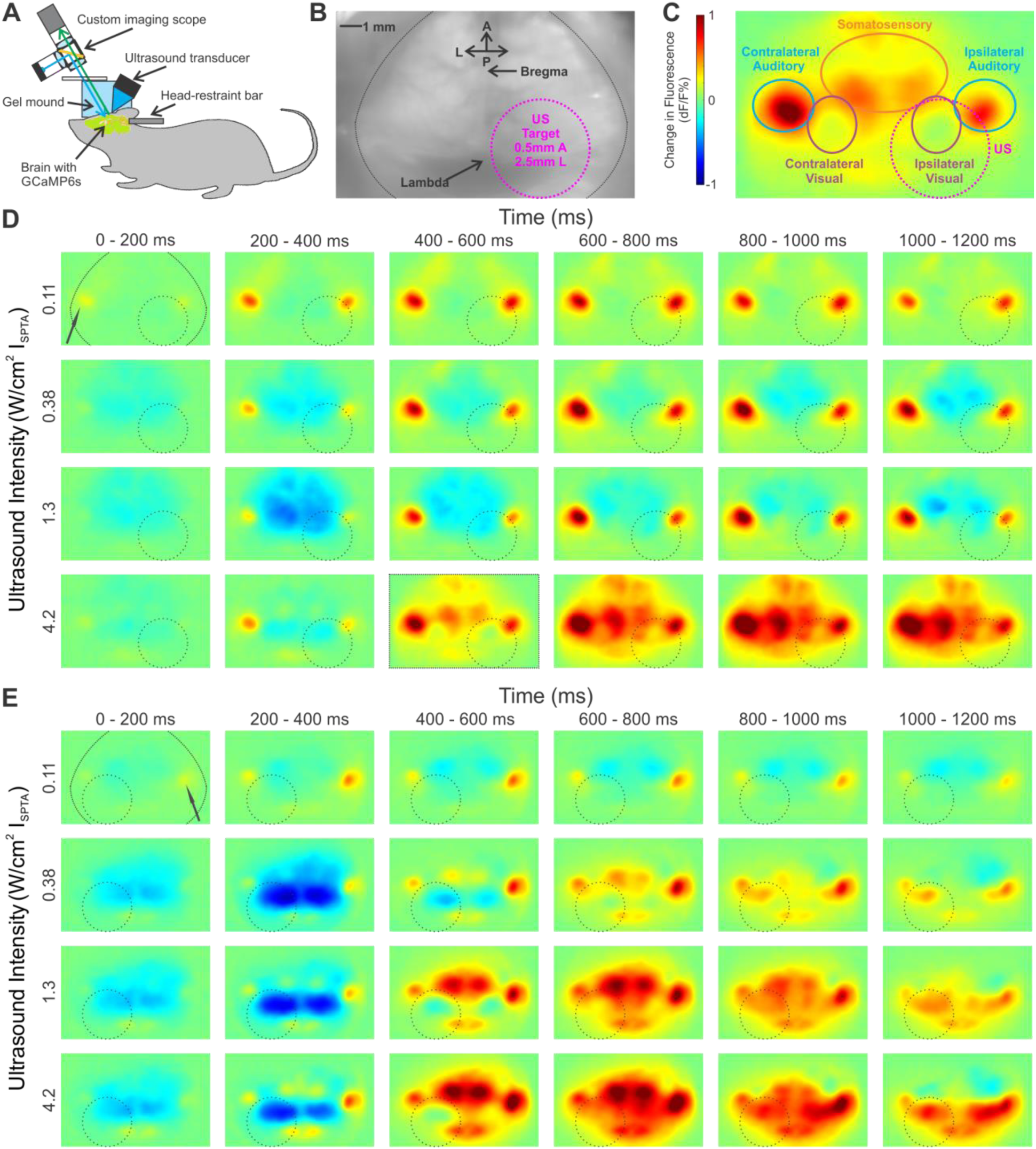
*Cortical responses to focused ultrasound*. **A**. Transgenic mice, with the genetically-encoded calcium sensor GCaMP6s expressed in the cortex, undergo a surgery where a metal head-restrained bar is implanted and their skull is thinned to obtain clear optical access to the brain. Ultrasound is delivered by a transducer that is held in a 3D-printed holder filled with ultrasound gel. To obtain both optical and ultrasonic access to the brain, a mound of clear ultrasound gel is used. The top surface is flattened with a glass plate to give clear optical images. **B**. Raw fluorescence image of a thinned-skull animal. The edge of the dorsal surface of the skull is covered by dental cement, along the black line at the edges. Anatomical landmarks such as the Bregma and Lambda sutures can be seen as well. **C**. A sample normalized change in fluorescence (dF/F) image, expressed in % change from baseline. The target region as well as outlines of the different sensory areas are shown. **D—E**. Responses of two representative animals to increasing intensities of ultrasound at different time points. The ultrasound target zone is shown as a dashed black circle. The contralateral auditory cortex is indicated with a black arrow. The approximate skull edge / dental cement outline is shown in the top left image. dF/F scale as in **C**.

The application of ultrasound to the visual cortex resulted in distinct and reproducible spatio-temporal patterns of cortical activation (Fig. 1, D-E). Surprisingly, the earliest regions to show a response were auditory cortices. At lower intensities of ultrasound, only the auditory cortices seemed to show an excitatory response reliably (top rows in Fig. 1, D-E), while other regions often showed a modest inhibitory signal or significantly delayed weak excitatory signal a few hundred milliseconds after ultrasound offset. At higher intensities, the auditory cortices showed excitatory signals early on (20–200 ms), during and immediately after the 80 ms ultrasound pulse, while other regions, including the visual cortex, became activated later, after around 400 ms (bottom rows in Fig. 1, D-E).

When we quantified the time course and strength of the calcium signals in the auditory cortex and the targeted visual cortex as a function of ultrasound intensity, we found that auditory regions were reliably activated earlier, and with lower powers of ultrasound, than the targeted visual cortical area (Fig. 2, A-B). Furthermore, the visual cortex targeted with ultrasound showed similar response kinetics and dependence on ultrasound intensity as the contralateral visual cortex, which was not targeted with ultrasound (Fig. 2, C-D). On both sides, the visual cortex showed an early fluorescence decrease, suggestive of cross-modal sensory inhibition (Iurilli et al., 2012).

**Figure 2.**
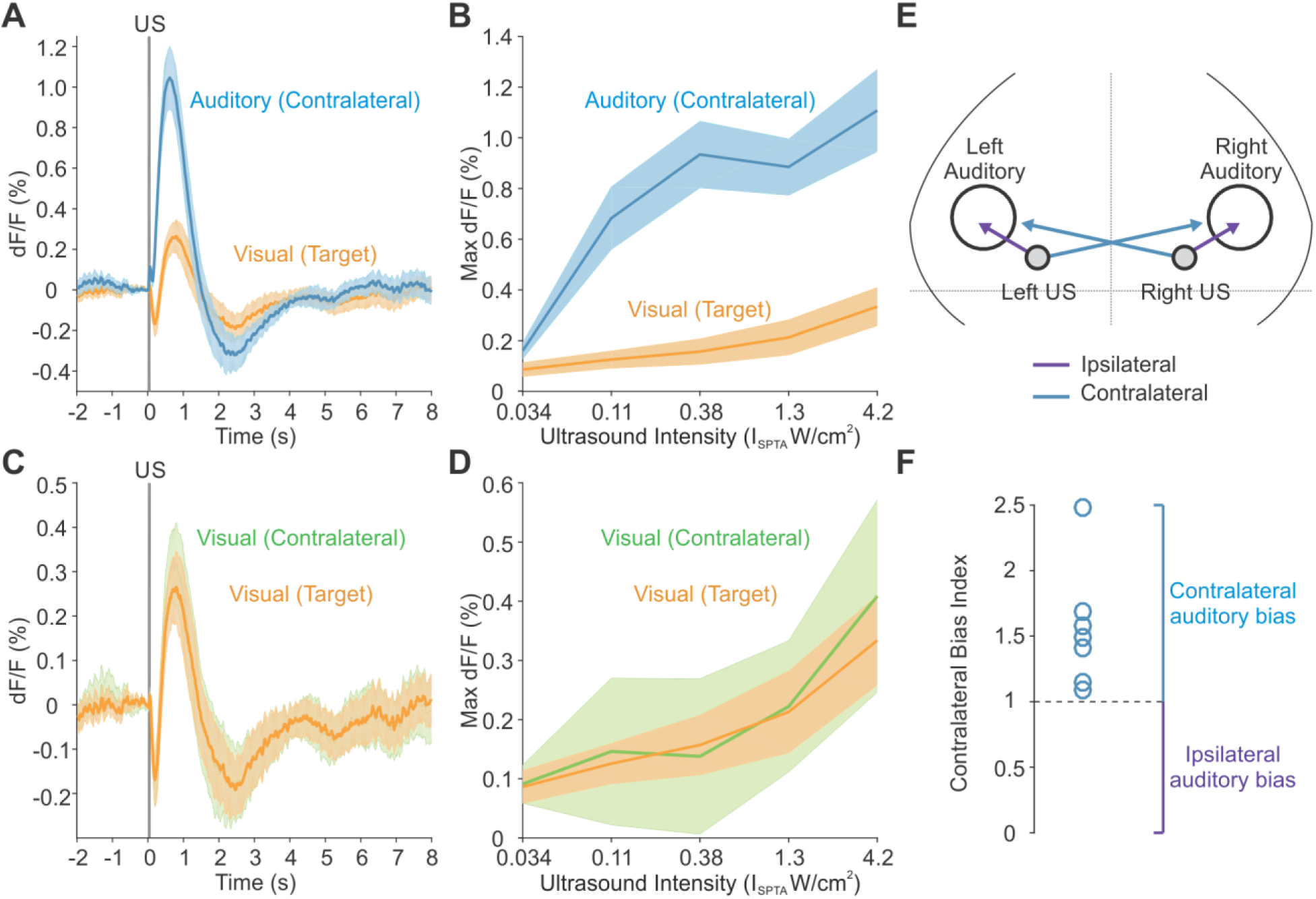
*Regional responses to ultrasound*. **A**. Response time course to ultrasound in the targeted region of visual cortex and the contralateral auditory cortex. **B**. Maximum dF/F signal in the targeted region of visual cortex and the contralateral auditory cortex in the first 2 sec after onset of ultrasound at different intensities. **C**. Response time course to ultrasound in the targeted region of visual cortex and the contralateral visual cortex. **D**. Maximum dF/F signal in the targeted region of visual cortex and the contralateral visual cortex in the first 2 sec after onset of ultrasound at different intensities. Mean traces in solid, SEM as shaded region (n=10 mice). **E**. Illustration of ipsilateral and contralateral relations between the ultrasound site and auditory cortex. **F**. Contralateral bias index for auditory activation with ultrasound in n=7 mice that were stimulated on both sides of the head.

The observation of strong and early signals in the auditory cortex led us to hypothesize that ultrasound was indirectly activating auditory pathways. To determine whether this activation was due to stimulation of inner ear structures or direct action on neurons in the auditory cortex, we compared auditory cortex activation ipsilateral and contralateral to the ultrasound focus. If the effects are mediated by the inner ear, one would expect the ear closest to the ultrasound focus to receive more of the stimulus, resulting in stronger activation of the contralateral auditory cortex due to auditory pathway decussation in the brainstem (Fig. 2E). Mice stimulated in separate trials at both right and left visual-cortical targets showed a clear contralateral bias, present in all animals tested (Fig. 2F), supporting the hypothesis that auditory cortex activation results primarily from effects on the ear closest to the ultrasound focus.

### Cortical response to ultrasound is similar to response to audible sound

To further elucidate the relative contributions of direct activation of the targeted region and indirect auditory effects on the spatiotemporal pattern of cortical activity elicited by UNM, we compared cortical responses to ultrasound, visible light flashes to the contralateral eye, and audible sound from a speaker driven at the same frequency as the ultrasound pulse repetition frequency of 1,500 Hz (Fig. 3A). Light flashes evoked a reproducible excitation of the visual cortex contralateral to the stimulated eye, typically followed by activation of the broader cortex (Fig. 3, B-D, top rows). In contrast, audible sound (108 dB) and ultrasound (I_SPTA_ 4.2 W/cm^2^) both induced strong activation of the contralateral auditory cortex, followed by a spreading change in activity to other cortical regions (Fig. 3, B-D, middle and bottom rows). Although some variability in the response pattern was observed across animals, responses to sound and ultrasound were always very similar within a specific animal.

**Figure 3.**
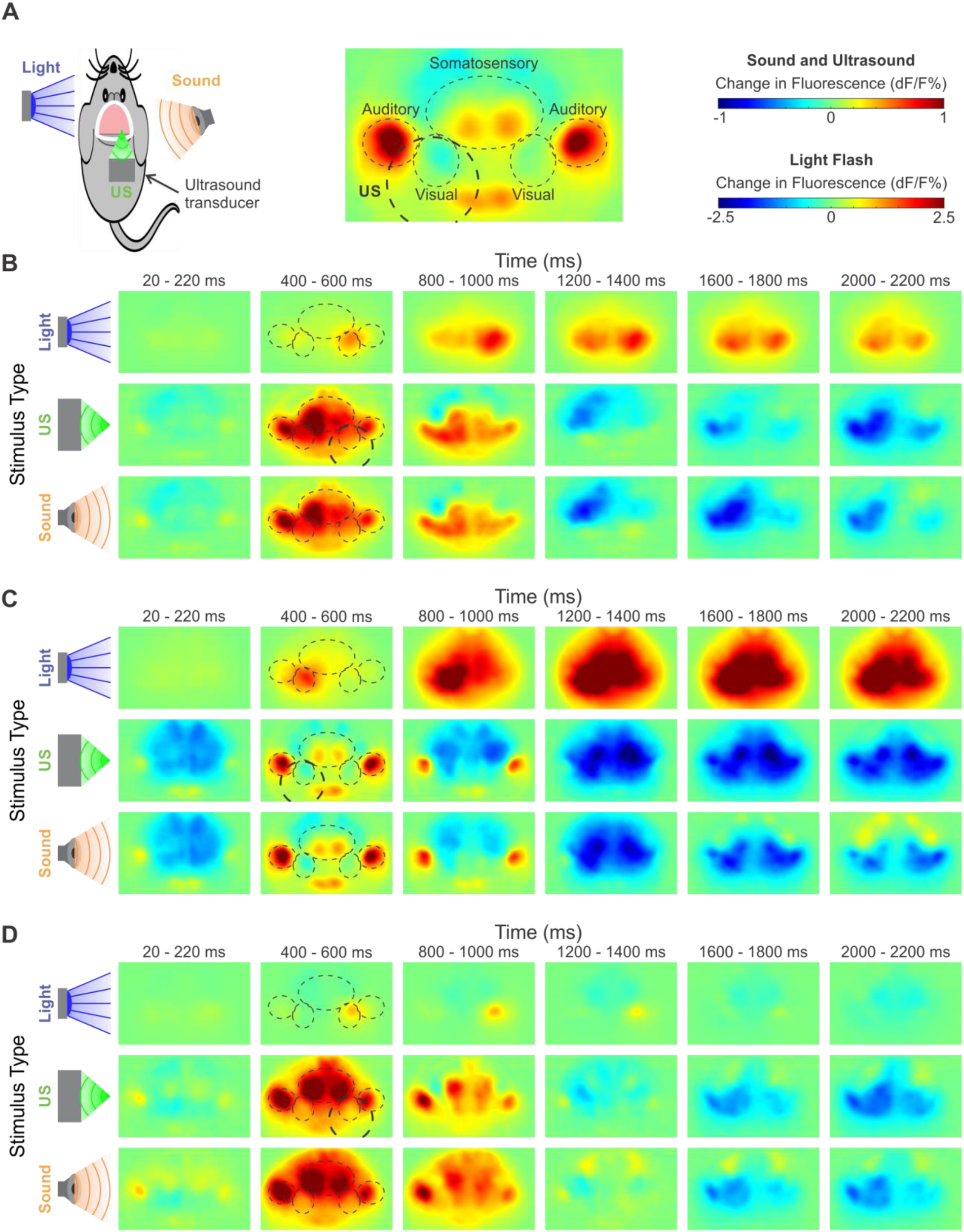
*Cortical responses to ultrasound, light flashes, and sound*. **A**. Diagram of experimental conditions and relevant cortical regions. **B-D**. Three representative cortical activation maps at different time points in response to light flashes to the contralateral eye, ultrasound (Ispta 4.2 W/cm^2^), and sound (108 dB). Relevant cortical regions are outlined to guide the eye. Ultrasound target indicated with thicker circle. dF/F scales as shown in **A**.

On average, the visual cortex responded most robustly to flashes of light to the contralateral eye, while showing a mixture of weaker inhibitory and excitatory responses to both sound and ultrasound, with similar time courses (Fig. 4A). By contrast, the contralateral auditory cortex displayed an immediate and robust signal to both sound and ultrasound (Fig. 4B), while showing a delayed and smaller positive signal in response to light flashes. To further quantify the similarity of brain activation patterns in time across the brain, we calculated a normalized similarity index between any two stimuli at a given time point (Fig. 4C) (see Methods for details). As expected, the two highest intensities of ultrasound (I_SPTA_ 4.2 and 1.4 W/cm^2^) had near-maximal similarity for the duration of imaging. More surprisingly, when the most intense ultrasound was compared to the most intense sound (108 dB), the spatiotemporal patterns were also highly similar at all time points. Meanwhile, light flashes induced a spatiotemporal signal pattern that was not only less similar, but had periods of negative similarity, indicating anticorrelated effects.

**Figure 4.**
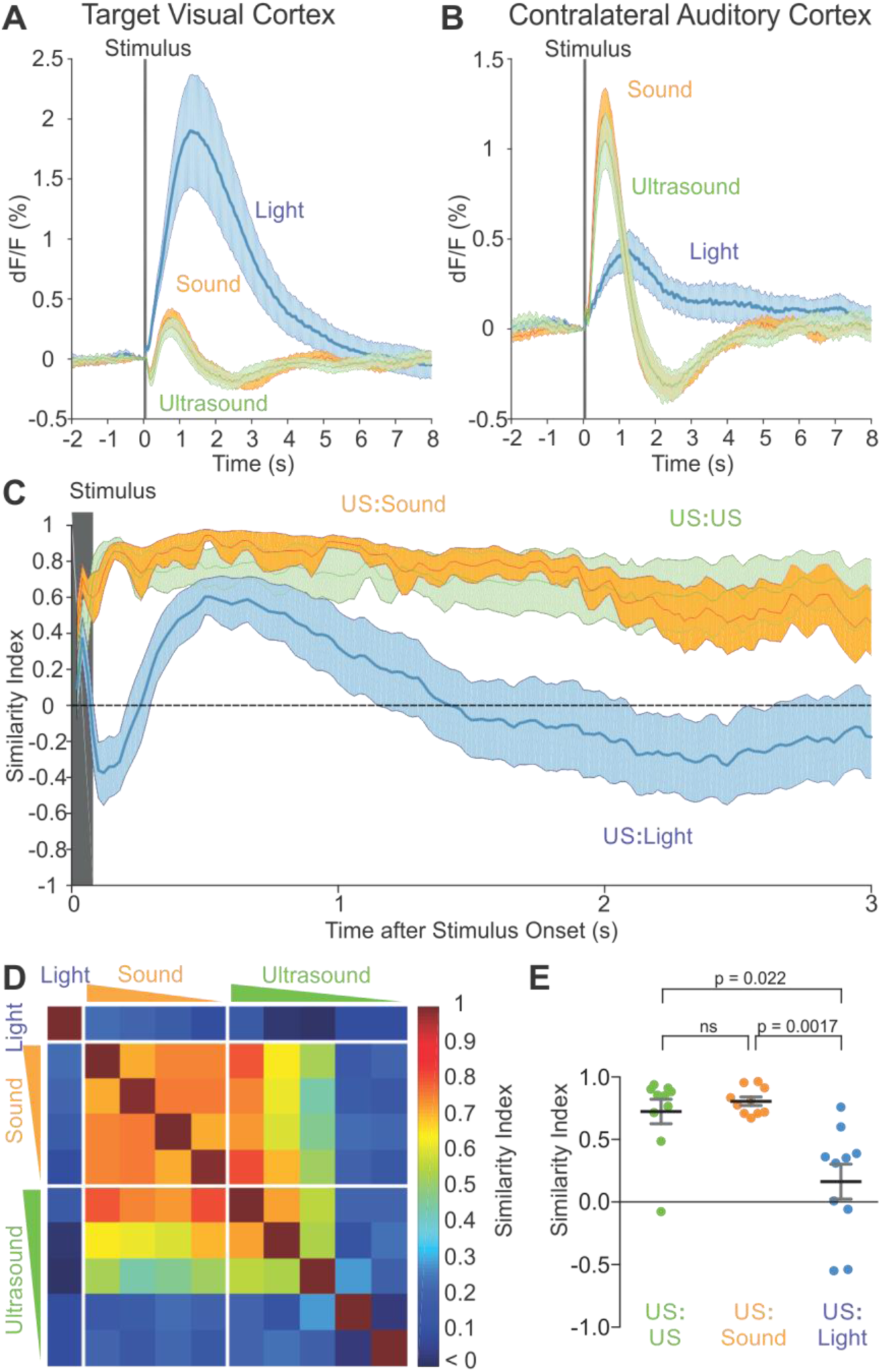
*Similarity of cortical responses to ultrasound, sound, and light flashes*. **A** Response time course of the ultrasound-targeted visual cortex to light, ultrasound, and sound. **B**. Response of contralateral auditory cortex to light, ultrasound, and sound. **C**. Spatial similarity index computed across time points for the indicated pairings of stimuli. Ultrasound at 4.2 W/cm^2^ was compared against ultrasound at 1.4 W/cm^2^, audible sound at 108 dB, and light flashes. **D**. Spatiotemporal similarity index computed over for the first 2 sec after stimulus onset across 10 animals as a matrix between all pairs of stimuli (10 in total). The 10 stimuli were: contralateral light flashes, four intensities of sound (decreasing from left to right or top to bottom), and five intensities of ultrasound (decreasing from left to right or top to bottom) as described in the Methods section. **E**. Statistical comparison of spatiotemporal similarity between the ultrasound, light, and sound conditions shown in **C**.

Expanding this analysis, we computed similarities among 4 intensities of sound, 5 intensities of ultrasound, as well as light flashes, over the first 2 s after stimulus onset, averaged across 10 animals (Fig. 4D). The spatiotemporal activity pattern induced by light flashes was dissimilar from all the other stimuli, while those induced by ultrasound and sound were similar to each other across several intensities. An analysis of the individual similarity indices in each of the 10 animals revealed significantly stronger correspondence between ultrasound and audible sound than between ultrasound and light flashes (Fig. 4E).

### Ultrasound and audible sound elicit movements consistent with startle reflex

Since most previous studies of UNM have used motor behavioral readouts, we asked whether limb movement elicited by ultrasound could be due to the secondary auditory effects identified in our imaging experiments. In particular, it is well known that unexpected sensory stimuli such as sound and air puffs can cause startle reflexes in animals, manifesting as movement (Galvani, 1970, Pilz, 2004, Vogel, 2005), and that strong stimuli can induce temporary arousal from anesthesia (Venes et al., 1971, Otto and Mally, 2003, March and Muir, 2005). To assess this possibility for ultrasound, we recorded electromyographic (EMG) signals from the left hindlimb as we applied UNM to the right visual cortex. This target area, located in the posterior region of the brain, has been shown by previous studies to be close to optimal for eliciting motor effects with ultrasound (Younan et al., 2013, Mehic et al., 2014, Ye et al., 2016, Kamimura et al., 2016). In addition to audible sound and ultrasound, air puffs to the face were used as a positive control for startle-eliciting stimuli, and light flashes were used to test whether strong visual activation could evoke movement. Strikingly, air puffs, audible sound and ultrasound all elicited similar EMG responses (Fig. 5A), suggesting their involvement in startle or arousal from anesthesia. In contrast, no motor responses were observed for light flashes, making it unlikely that ultrasound causes a startle reflex by generating a phosphene. This result is in line with the lack of observed direct activation of the visual cortex by ultrasound.

**Figure 5.**
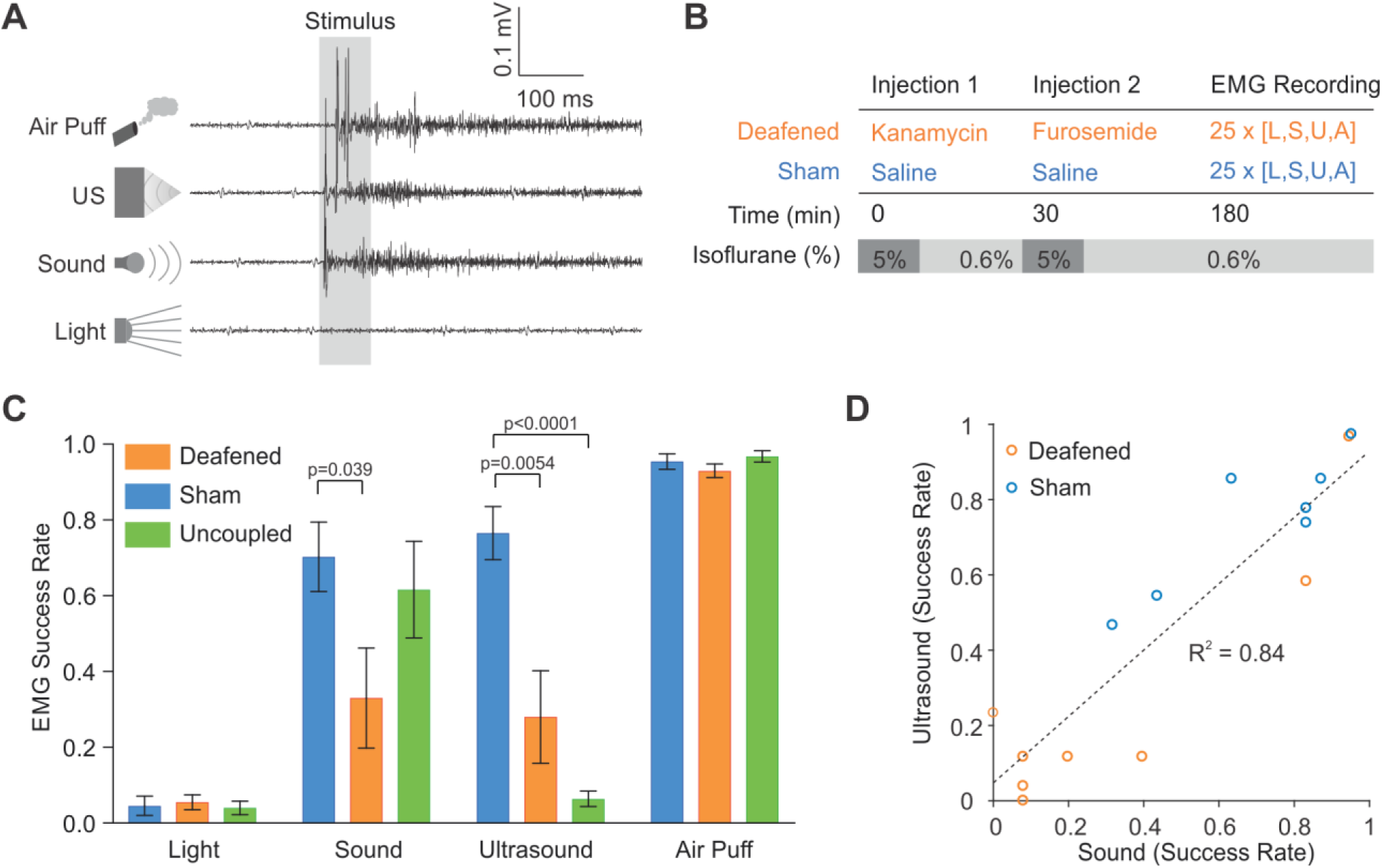
*Motor responses to ultrasound, sound, light, and air puffs*. **A**. Representative EMG recordings from mice in response to the four indicated stimuli. Ultrasound and sound were at 4.2 W/cm^2^ and 108 dB, respectively. **B**. Protocol for mouse deafening. **C**. Stimuli responses of chemically-deafened animals (n=8), saline injected animals (n=7), and “no gel” animals (n=5). **D**. Correlation in response rates to ultrasound and sound across individual animals included in the sham and deafened groups.

### Chemical deafening reduces motor responses to ultrasound and audible sound

The fact that sound and ultrasound both elicited similar EMG signals suggested that the motor effects of UNM may, at least partially, be due to auditory-mediated startle rather than direct effects of ultrasound on the motor cortex. To further evaluate this possibility, we chemically deafened a subset of animals using a cocktail of kanamycin and furosemide (30 min later) (Fig. 5B) (Taylor et al., 2008, Oesterle et al., 2008). This cocktail is expected to produce partial deafening within 30 minutes after furosemide administration (Li et al., 2011, Xia et al., 2014). Strikingly, chemical deafening greatly reduced the motor responses to both sound and ultrasound, while leaving air puff response rates unaltered (Fig. 5C).

In an additional experiment, we tested uncoupling the ultrasound transducer from the head by not using ultrasound gel. This resulted in a near complete abolishment of motor responses to ultrasound in animals that still responded to sound. This suggests that the auditory activation from ultrasound requires contact, and is not caused by airborne transmission of sound waves from the transducer to the ears.

As expected for the chemical deafening protocol, animals had variable hearing loss, and saline-injected sham animals also showed some variability in their response to auditory stimuli, possibly due to differences in sensitivity to anesthetics or tendency for startle. This allowed us to examine the correlation between each animal’s responsivity to sound and to ultrasound (Fig. 5D). The strong correlation between responses to these two stimuli (R^2^ = 0.84), is further evidence of the involvement of auditory pathways in ultrasound-induced motor responses.

In addition to the EMG results above, we attempted to observe the impact of chemical deafening on cortical calcium signals. However, chemical deafening in the older transgenic animals we used for imaging resulted in high mortality. We suspect that this is due to the age of the mice, as chemical deafening has not been tested in older animals. Unfortunately, the surgery to implant a head plate and thin the skull necessitates older animals (20+ weeks) for good post-surgical recovery.

## Discussion

Our study reveals that focused ultrasound applied to a non-auditory brain region in mice produces strong activation of auditory cortex and additional brain regions with spatiotemporal dynamics closely resembling those elicited by audible sound. This activation is sufficient to produce motor behavior consistent with an auditory startle reflex, occurring via pathways involving the inner ear, as documented by the inhibitory effects of chemical deafening. Compared to the robust activation of the visual cortex with light flash stimuli, no direct activation of this cortical region with ultrasound was observed.

The precise mechanisms by which ultrasound at 500 kHz, a frequency normally inaudible to animals such as mice and humans, activates auditory pathways, is an important topic for future study. The mechanisms by which air-coupled ultrasound (Westervelt, 1963, Yoneyama et al., 1983, Averkiou et al., 1993) and soft-tissue conducted sound (Wever and Bray, 1937, Goodhill and Holcomb, 1955, Mauldin and Jerger, 1979, Dobrev et al., 2017) activate the auditory system are relatively well understood. It is also known that ultrasound can elicit auditory sensations in humans when coupled through bone (Pumphrey, 1950, Pumphrey, 1951, Deatherage et al., 1954, Corso, 1963). However, there is still no consensus on how soft-tissue coupled ultrasound activates the auditory system (Deatherage et al., 1954, Haeff and Knox, 1963, Dieroff and Ertel, 1975, Foster and Wiederhold, 1978, Gavrilov, 1984, Lenhardt et al., 1991, Dobie and Wiederhold, 1992, Magee and Davies, 1993, Hosoi et al., 1998, Nishimura et al., 2003). Potential mechanisms include mode conversion between primary compressive ultrasound waves and shear waves within bone and the brain’s soft tissue (Clement et al., 2004, White et al., 2006, Vignon et al., 2010, Gennisson et al., 2013), leading to mechanical activation of ear structures. These secondary waves would have primary frequencies determined by the pulse repetition frequency and the mechanical properties of tissue, as well as a broadband component arising at the start and end of each tone burst, likely within the audible range of 1 to 100 kHz (in mice). The companion paper by Guo et al. (https://doi.org/10.1101/233189) suggests that the auditory coupling involves cochlear fluids.

The use of mice as a model allowed us to take advantage of the availability of transgenic animals expressing a cortex-wide fluorescent reporter of calcium. However, it is possible that the small size of the mouse head makes these animals particularly susceptible to the auditory side-effects of ultrasound, and that skull reflections at this scale could generate standing waves leading to more complex pressure patterns and mechanical forces (O’Reilly et al., 2010, Younan et al., 2013). These concerns are mitigated by the corroborating findings of Guo et al. (https://doi.org/10.1101/233189) in the accompanying study, which used guinea pigs with brain volumes 8 times larger than in mice. The ability of ultrasound to elicit audible sensations in humans has also been reported in studies dating back to 1950 (Pumphrey, 1950). Nevertheless, further experiments in animals with larger head sizes are needed to assess the extent of ultrasound-induced auditory effects across species.

Motor responses to ultrasound, as well as those caused by audible sound and air puffs, may depend on the depth of anesthesia. In previous UNM studies, isoflurane has been used at levels between 0.02 and 0.6% (King et al., 2013, King et al., 2014, Ye et al., 2016), while deeper anesthesia made it difficult to obtain motor responses (King et al., 2013). For studies utilizing isoflurane, a key factor implicated in UNM efficacy was light-anesthetic conditions where the animal exhibited spontaneous movement as assessed by EMG signals (King et al., 2013, King et al., 2014, Ye et al., 2016). Depending on the body temperature of the mouse, anesthesia at 0.5 to 1.5% is the range in which animals begin to lose reflexes including those to noxious stimuli such as tail pinches (Werner et al., 2011). The ketamine-xylazine cocktail, also used in UNM studies (Tufail et al., 2010, Yoo et al., 2011, Kim et al., 2014, Younan et al., 2013, Mehic et al., 2014), results in variable anesthetic depth due to its short half-life, and it is unclear what level of anesthesia animals experienced when motor responses were measured (Tufail et al., 2010, Yoo et al., 2011, Kim et al., 2014). Some papers specifically state that animals retained the tail-pinch reflex during their experiments, suggesting light anesthesia levels (Mehic et al., 2014).

Using ultrasound parameters consistent with previous UNM studies (Tufail et al., 2010, King et al., 2013, Mehic et al., 2014), we were unable to obtain evidence of direct neuromodulation at the targeted cortical region. Although this region exhibited reproducible inhibition and activation, this was part of a larger activity pattern encompassing multiple brain regions, with an almost identical response in the symmetric contralateral cortex. This activation pattern did not resemble the activity evoked by the cognate light flash sensory stimulation, while air-coupled sound created nearly identical spatiotemporal activity patterns as ultrasound. These patterns were consistent with previous literature on cross-modal sensory connectivity (Iurilli et al., 2012) (Fig. 6).

**Figure 6.**
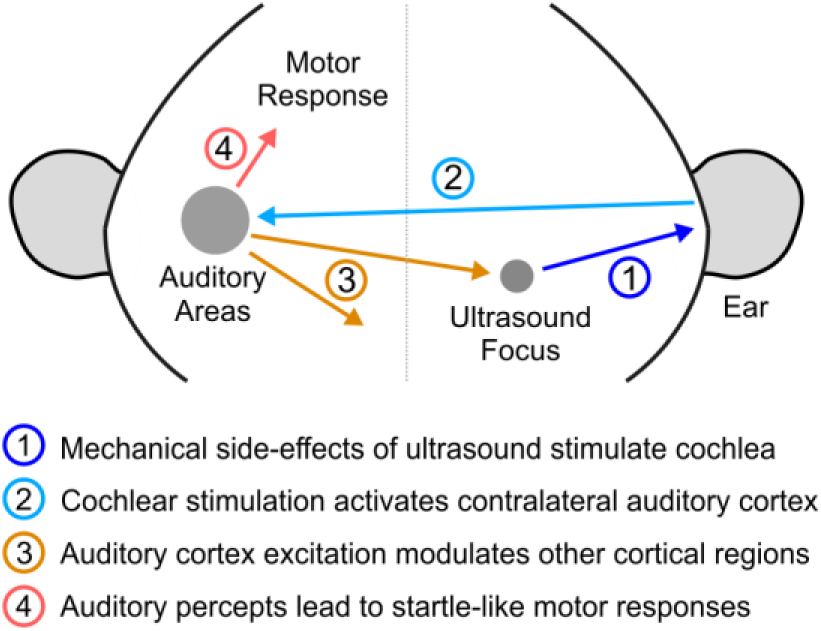
*Illustration of indirect auditory effects of ultrasonic neuromodulation*. 1. Ultrasound application leads to mechanical waves stimulating the inner ear structures of the cochlea. 2. The activation of the cochlea leads to excitation of auditory pathways, including the contralateral auditory cortex. 3. Cross-modal projections from these auditory regions lead to modulation of neural activity across the cortex, including the neurons that are within the focal zone of ultrasound. The timing and sign of this modulation is nearly identical to that caused by air-coupled sound. 4. The auditory percept can also lead to startle-like motor responses.

However, we caution that our results do not conclusively demonstrate that ultrasound is unable to produce direct neuromodulatory effects. For example, if ultrasound activates subcortical regions, we would not have been able to observe this with wide-field fluorescence. Likewise, modulation of finer aspects of neuronal excitability, such as synaptic vesicle release, action potential time course, or magnitude and duration of evoked potentials, could have been difficult to detect in our experiments. It is also possible that alternative ultrasound parameters not tested in this study, including potentially higher stimulus intensities, could produce direct modulation. Furthermore, our findings of off-target auditory effects do not, in our view, disqualify UNM from serving a useful function in neuroscience and clinical applications. Other widely used technologies such as transcranial magnetic stimulation (TMS) are also known to produce sensory side-effects, which can be accounted for with appropriate sham controls.

Finally, stand-alone UNM is only one of several methods through which transcranial ultrasound can be used to affect brain function, others including targeted ablation (Fry, 1977, Fry and Goss, 1980, Huang and Hynynen, 2011, Lipsman et al., 2013, Chang et al., 2015, Arvanitis et al., 2016, Elias et al., 2016); localized blood-brain barrier opening (Hynynen et al., 2003) to directly affect local neural activity (Chu et al., 2015, Downs et al., 2017) or to deliver genes, proteins, cells, and small molecules (Kinoshita et al., 2006a, Kinoshita et al., 2006b, Treat et al., 2007, Choi et al., 2010, Choi et al., 2011, Chen et al., 2013a, Chen et al., 2014, Wang et al., 2015, McDannold et al., 2015, Airan et al., 2017a, Airan et al., 2017b); and emerging sonogenetic approaches (Heureaux et al., 2014, Ibsen et al., 2015). These methods can all leverage the technological developments facilitating noninvasive focal ultrasound delivery to the brain (Fan and Hynynen, 1994, Hynynen and Jolesz, 1998, Sun and Hynynen, 1998, Tanter et al., 1998, Tanter et al., 2000, Pernot et al., 2003), theoretically requiring only an intravenous injection as the most invasive step. Notwithstanding these important caveats, it is clear that further investigation of both direct and indirect effects of ultrasound, and the development of proper sham controls for UNM, will be critical for the future of this field.

## METHODS

### Animal Use

For this study, mice were used in accordance with animal procedures approved by the Institutional Animal Care and Use Committee at the California Institute of Technology. For imaging studies, transgenic male mice, C57BL/6J-Tg(Thy1-GCaMP6s)GP4.12Dkim/J (Thy1-GCaMP6s) GP4.12Dkim (The Jackson Laboratory, Stock No. 025776), over 20 weeks of age and weighing over 35g were used. Due to the very large surgery area to expose the bilateral auditory regions, only these large animals could undergo surgery and remain healthy. For EMG experiments, male C57BL/6J mice, 8-12 weeks of age, weight of 25-30 g, were used. This age group and size more closely matches previous studies of motor responses to ultrasound and have better health outcomes in response to chemical deafening procedures.

### Animal Surgery

Anesthesia was induced by placing mice in a clean induction chamber and delivering 5% isoflurane. The animals were then placed in a stereotax and the head was held steady using ear bars and a nose cone. Anesthesia was maintained via delivery of isoflurane (1.5~2%) through the nose cone. Body temperature was maintained using a heating pad. Extra care was made to ensure the eyes remained protected using ophthalamic ointment. Briefly, fur was removed using hair removal cream and the exposed scalp sterilized using chlorhexidine. The skull was then exposed via an incision along the midline and laterally above the cerebellum. For mice used in imaging, a skull-thinning procedure was performed (Grutzendler et al., 2002, Yang et al., 2010). A micro-burr bit (19007-07, Fine Science Tools, Inc., Foster City, CA) was used to gently thin the skull while cooled saline was used to prevent any thermal buildup from the procedure. The thin-skull procedure was chosen for a number of reasons. First, thin-skull surgeries (as opposed to craniotomies) do not lead to significant changes in the brain due to the surgeries, such as spine turnover and glial buildup (Xu et al., 2007). Second, craniotomies are typically sealed with glass coverslips, which will create an acoustically-mismatched surface. Mouse skulls have been shown to be transparent to 500 kHz ultrasound (King et al., 2013) and in particular, skull-thinning to optical clarity reduces skull thickness to ~ 50 microns, further lessening any aberration effects on the ultrasound field. Finally, craniotomies to expose the area of cortex that was imaged in this study carry higher risks both during surgery and recovery. Mice used for EMG did not undergo the skull-thinning procedure as optical access to their brain was not needed, and previous studies have demonstrated that these skulls are acoustically transparent to 500 kHz ultrasound (King et al., 2013) . All animals were then implanted with a stainless steel head-restraint plate using dental acrylic (C&B-METABOND, Parkell, Inc., Edgewood, NY). The exposed skull was covered using quick-acting silicone (Kwik-Sil, World Precision Instruments, Inc. Sarasota, FL) to form an easily removable silicone plug. Animals were then placed in a heated clean cage and allowed to recover.

### Experimental Preparation

Each experiment day, anesthesia was induced by placing mice in a clean induction chamber and delivering 5% isoflurane. As soon as voluntary movement ceased, mice were quickly moved to the head-restraint setup and maintained at 2% isoflurane for preparation. The silicone plug was removed. A 3D printed well was attached to the dental acrylic well on the skull and to the head-restraining bars. The ultrasound transducer, angled at 60 degrees from parallel was brought to the approximate region using a 3D-printed piece that clipped onto the transducer holder and allowed targeting. This piece was then removed to allow optical access to the focus. A fiber-optic hydrophone was then brought to the target location. The well was then filled with ultrasound gel. For imaging, a glass plate was brought down to flatten the top surface so that imaging can be performed through the gel. Air bubbles were removed using a syringe. To keep the anesthesia protocol as similar as possible to other studies utilizing isoflurane (Ye et al., 2016), anesthesia was maintained at 2% for a total of 34 min. Anesthesia was then reduced to 0.5% for 5~10 min as needed for tail-pinch reflexes to return, and then increased to 0.6%. The ultrasound transducer was then adjusted using a 4-axis micrometer (XYZ + axial) to maximize the pressure at the hydrophone using short, low-intensity pulses (50 us pulses, 500ms between pulses, 100 kPa peak pressure). Experiments were then started.

### Experimental Design for Imaging

All imaging animals except for those used in Fig. 2E and F underwent 200 blocks of experiments. In each block, a trial of each stimulus (light flash to the eye contralateral to the ultrasound target, 5 intensities of ultrasound, 4 intensities of sound) was presented once in random order. Data for Fig. 2E and F were obtained using 2 blocks of experiments, one where ultrasound was targeted to the left target coordinate, and another targeting the right coordinate, both using only ultrasound with I_SPTA_ of 4.2 W/cm^2^. All ultrasound stimuli were of 500 kHz acoustic frequency, 100 cycles/pulse, 1.5 kHz pulse repetition frequency, and 120 pulses. This yields a duty cycle of 30% and stimulus duration of approximately 80 ms. The highest intensity of ultrasound had an Ispta of 4.2 W/cm^2^. This value was chosen to correspond to the value determined by King et al (King et al., 2013) to be the range at which UNM becomes reliable. Further studies in mice corroborate that this Ispta is at or higher than levels needed for neuromodulation (Tufail et al., 2010, Mehic et al., 2014). The timing parameters were also chosen to mimic those tested by these studies of motor responses in mice (Tufail et al., 2010, King et al., 2013, Mehic et al., 2014). Indeed, the highest intensity ultrasound reliably elicited motor responses in test mice as well, confirming the suitability of this intensity value. The lower intensities of ultrasound were generated by reducing the voltage sent to the RF amplified such that each subsequent ultrasound waveform had 30% intensity of the previous intensity, namely Isptas of 1.3, 0.38. 0.11, and 0.034 W/cm^2^. The sound intensity for the loudest stimulus was adjusted by changing the driving voltage such that the auditory response in the contralateral cortex was similar to that evoked by ultrasound. The waveform for the loudest sound were created using 120 +5 V pulses of 200 us duration at 1.5 kHz (30% duty cycle). The subsequent intensities were created by reducing the duty cycle by 30%, thus reducing electrical power input by 30%, but holding driving voltage constant. Thus, 9%, 2.7%, and 0.81% duty cycle waveforms were used. The light flash intensity was chosen so that the maximal cortical activation in the first 500 ms was of similar magnitude as that elicited by the strongest sound and ultrasound intensities.

### Experimental Design for Electromyography

To chemically deafen animals, at the start of the preparation period, an injection of kanamycin (1g/kg SC, K0254, Millipore Sigma, Inc., St. Louis, MO) was given. 30 minutes later, Furosemide (200mg/kg IP, PART NUM) and saline (1.5 mL SC) were given. For saline control animals, the 3 injections were all done using 0.9% saline solution using the same timing and anesthetic doses. For the gel-uncoupled controls, no ultrasound gel was used between the transducer and the skull; all 3 injections again were with 0.9% saline solution and identical anesthetic regimen. Thirty minutes after the final injection, 25 blocks of one trial each of sound, ultrasound, air puff, and light flashes were obtained with an ITI of 10 sec. At 150 minutes after the final injection, another 25 trials were obtained using the same stimuli and ITI. This last block was used for analysis. All four stimuli were randomly ordered within a block.

### Experimental Control

Experiments were controlled by custom software, written in LabVIEW (National Instruments, Austin, TX). A PXIe chassis (PXIe-1073), housing a data acquisition board (DAQ) (PXIe-6363), and a function generator (FxnGen) (PXI-5421), all from National Instruments (National Instruments, Austin, TX), was used to interface with other hardware and circuits (detailed below).

### Ultrasound Generation, Calibration, and Delivery

A 500 khz ultrasound transducer (AT24020, Blatek, Inc., State College, PA), with focal distance 30 mm and focal diameter 4.4 mm was used in all experiments. A timing counter on the DAQ board was used to generate a set number (NP) of trigger pulses at the PRF that was sent to the FxnGen to generate a set number (NC) of cycles of ultrasound at the acoustic frequency. This signal was amplified using an RF amplifier (240L, Electronics & Innovation, Ltd, Rochester, NY), and the amplified output signal was used to drive the ultrasound transducer. The transducer was housed in a 3D printed holder for experiments. Calibration was done with a fiber-optic hydrophone system (FOH, Precision Acoustics, Dorchester, UK) using hydrophones (PFS and TFS, Precision Acoustics, Dorchester, UK) calibrated at the National Physical Laboratory (London, UK).

### Non-Ultrasonic Stimuli Generation and Delivery

Three non-ultrasonic stimuli: light flashes, air-coupled sound, and air puffs, were used as well. The timing signal for non-ultrasonic stimuli was generated using two timing/counter channels on the DAQ board. This signal was passed through an optoisolator (HCPL2630M, ON Semiconductor, Phoenix, AZ). Using AND gates (CD74AC08E, Texas Instruments, Dallas, TX) and three digital logic output channels on the DAQ (one for each modality), this timing signal was routed to three independent NMOS circuits to power an LED, speaker, or solenoid valve at their appropriate driving voltages. Sound was generated by delivering a timed +5V pulse that drove a speaker (SP-1813-2, Soberton, Inc, Minneapolis, MN) placed near the mouse’s ear. The waveform for the loudest sound were generated by a train of 200 us-long +5V pulses at a repetition frequency of 1500 Hz for a total of 120 repeats, resulting in a 30% duty cycle waveform that was approximately 80 ms in duration. Lower intensity sounds were generated by reducing the duty cycles by 30% successively, but with the same driving voltage, frequency, and number of pulses. The speaker volumes were measured to be 108 dB, 98.8 dB, 85.3 dB, and 69 dB for the powers used. Light flashes were generated by delivering timed +4V pulses to an LED (SP-01-B6, Quadica Developments, Inc, Alberta, CA) coupled to a flexible plastic optic fiber (02-551, Edmund Optics, Inc, Barrington, NJ) that was brought to the animal’s eye. For light flashes, the stimulus duration was 19 ms in order to keep artifacts due to increased light limited to a single imaging frame. Solenoid valves (RSC-2-12V, Electric Solenoid Valves, Islandia, NY) for air puff stimulation were driven by a 80 ms-long +12V pulse. Electrical power for each was supplied by independent benchtop power supplies (1621A, BK Precision Corp, Yorba Linda, CA).

### Image Acquisition and Analysis

Imaging was performed using a home-built optical scope with 1.42x minification (The objective had focal length 60 mm, AC254-060-A, Thorlabs, Inc. Newton, NJ. The tube lens had focal length 40mm, AC254-040-A, Thorlabs, Inc. Newton, NJ. The lenses were adjusted so that the field of view at the focus was 16 mm x 10 mm). Images were collected at 50 Hz using a camera (GS3-U3-23S6M-C, FLIR Systems, La Mirada, CA). The exposure signal from the camera was used as the master timing signal to initiate trials. Fluorescence excitation light was generated by a 470 nm LED light source (SP-08-B6, Quadica Developments, Inc, Alberta, CA) powered by a benchtop power supply (1621A, BK Precision Corp, Yorba Linda, CA). A fluorescence filter set suitable for GCaMP imaging was used (excitation/dichroic/emission filter set GFP-4050B-000, Semrock, Inc. Rochester, NY). Image analysis was performed using custom code written in MATLAB. Each image frame was spatially filtered with a 500 um square filter to reduce noise. Temporal averaging was only used for creating maps of normalized changes in fluorescence (dF/F), and were avoided otherwise to maintain temporal fidelity of neural responses. In images of light flash trials, the first frame contained optical contamination from the LED flash, and thus the dF/F for this frame was set to 0 in plots of time courses. This was not done for images in response to other stimulus modalities. The first frame was excluded from analyses of spatiotemporal similarity and in generating the dF/F maps where noted in figures. Negative dF/F was interpreted as inhibitory signal (Akerboom et al., 2012, Tecuapetla et al., 2014, Kim et al., 2016, Mazo et al., 2016, Wang et al., 2017). The ROI for the target was chosen as shown in Fig. 1B. The contralateral control was chosen as the mirror-symmetric region (2.5 mm lateral and 0.5 mm anterior of Lambda) on the other side of midline. Sensory cortices were identified based on a stereotactic atlas (Franklin and Paxinos, 2013) with further refinement from functional imaging studies (Guo et al., 2012, Garrett et al., 2014, Issa et al., 2014, Tsukano et al., 2016, Juavinett et al., 2017). The locations of the visual and auditory cortices were further confirmed by their activation to visual and auditory stimuli, respectively. The ROI coordinates for the auditory cortex were chosen to be the location with maximal activation to sound.

### Contralateral Bias Index

In order to assess the amount of contralateral auditory cortex activation in contrast to the ipsilateral auditory cortex, we defined a contralateral bias index that analyzed the simultaneous bilateral auditory cortex responses when ultrasound was applied first to the left target site and then the right target site, or vice versa. This index was designed to account for potential signal imbalances between the two cortices, hearing differences in ears, or animal responsivity during the two blocks of stimulation (left and right targets). We defined the contralateral bias index as follows.

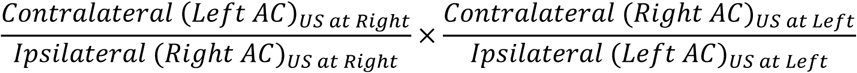

### Similarity Index

To analyze the spatiotemporal similarity of cortical responses to two stimuli, we defined a similarity index between two stimuli at any given time point. To do this, at each time point, for pixels corresponding to visible cortex, we took the sum of the pixel-by-pixel product of dF/F for the two stimuli and divided it by the square root of the sum of (dF/F)^2^ for each stimulus.

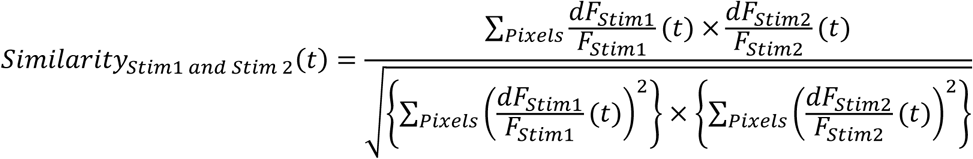

Thus, any spatiotemporal map that was identical but different in magnitude would have a similarity of 1. This index can be thought of as a non-mean-subtracted correlation of pixels in time. Standard metrics of correlation subtract by a population mean; however, our values were already baseline subtracted for fluorescence and further subtraction can alter the polarity (excitatory vs inhibitory) of the dF/F signal. As mentioned before, the similarity index analysis starts at the second frame after stimulus onset to avoid optical contamination in the light flash case.

### Electromyography Acquisition and Analysis

EMG signals were acquired using subdermal needles (RLSND110-1.0, Rhythmlink LLC, Columbia, SC) inserted into the left hindlimb. Reference and ground leads were placed in the scruff of the skin on the back. EMG signals were amplified using an extracellular amplifier (Model 1800, A-M Systems Inc, Sequim, WA) using a 100 Hz high-pass filter, a 5000 Hz low-pass filter, and 60 Hz notch filter and recorded by the analog input channels on the DAQ. Data was analyzed using custom code written in MATLAB (Mathworks, Natcik, MA). First, the DC offset of the EMG signal was subtracted out by using the mean of the prestimulus period. This signal was then rectified. This rectified signal was then smoothed using a bilaterally truncated Gaussian filter with width of 40 ms and full-width half-max of 10 ms. We then calculated the ratio between the average of this signal in the 150 ms time period between 80 (just after stimulus offset) and 230 ms after stimulus onset and the 150 ms preceding stimulus onset. A cutoff ratio of 1.25 was used to determine if a motor response had occurred. The time period was chosen to increase sensitivity to both shorter and longer contractions by taking into consideration of latencies to motor responses and contraction durations. In addition, by sampling the time period after offset of electrical currents to drive stimuli, it avoids any potential contamination of EMG signals by electrical interference, a possibility which was noted in one study (Younan et al., 2013).

## Acknowledgements

The authors thank Hubert Lim, Hongsun Guo and Sangjin Yoo for helpful discussions and input on the manuscript, and members of the Shapiro and Tsao labs for assistance with experiments. This research was supported by NIH BRAIN Initiative grant R24MH106107 (Co-PIs D.Y.T. and M.G.S.). Related research in the Shapiro Laboratory is also supported by the Heritage Medical Research Institute and the Packard Fellowship in Science and Engineering.

## Author Contributions

T.S., M.G.S. and D.Y.T. conceived the study. T.S. designed and performed all experiments and analyzed the data. T.S., M.G.S. and D.Y.T. interpreted the results and wrote the manuscript.

